# The genetic architecture of shoot and root trait divergence between upland and lowland ecotypes of a perennial grass

**DOI:** 10.1101/301531

**Authors:** Albina Khasanova, John T. Lovell, Jason Bonnette, Jerry Jenkins, Yuko Yoshinaga, Jeremy Schmutz, Thomas E. Juenger

## Introduction

Adaptation to abiotic stress is an important driver of contemporary evolution in plant populations. Abiotic stressors have been implicated as driving factors in ecological speciation (Stebbins, 1952;Lexer & Fay, 2005), where populations have diverged across a number of traits, exhibit different niche characteristics, and eventually become reproductively isolated (Clausen, 1951; Lowry, 2012; Yardeni *et al.*, 2016). Local adaptation to soil water availability is an especially important driver of plant evolution (Stebbins, 1952; Rajakaruna, 2004; Kooyers *et al.*, 2015) and can impose strong natural selection on plant populations that can lead to the formation of ecotypes that are differentially adapted to xeric and mesic habitats (Porter, 1966; Joly *et al.*, 1989; Kumar *et al.*, 2008). Xeric and mesic ecotypes are often characterized by the divergence of common suites of morphological and phenological traits (Clausen, 1951; Lowry, 2012) related to maintaining water status and avoiding periods of drought (Chapin *et al.*, 1993; Markesteijn & Poorter, 2009; Juenger, 2013).

Although leaf and shoot traits are important drivers of adaptation to drought (Tsialtas *et al.*, 2004; Carmo-Silva *et al.*, 2009; Juenger, 2013), the properties of root systems determine plant water access and can place constraints on shoot water status (Price et *al*., 2002; Hund *et al.*, 2009). Shoot traits may be related to root traits through genetic correlation (Bouteille *et* al., 2012) or dependent upon root traits though resource allocation tradeoffs (Hammer *et al.*, 2009), including changes in carbon allocation between root and shoot systems (Hummel *et al.*, 2010). Higher root mass ratio (RMR) increase water foraging capability to maintain plant water status, which can be accomplished by allocating more resources towards roots (Knights *et al.*, 2006) or by inhibiting above ground growth (Hendricks *et* al., 2016). Many root and shoot traits show correlated responses to water limitation and varied degrees of morphological and physiological integration. For example, decreased soil moisture in xeric environments is positively associated with high specific leaf area (SLA; Ramírez-Valiente & Cavender-Bares, 2017) and high specific root length (SRL; Comas *et al*, 2013) while increased soil moisture is positively associated with low SLA and low SRL (Price *et al.*, 2017). Increased SRL can increase plant water acquisition under drought without increasing carbon allocation per root length by producing longer and thinner roots (Comas *et al.*, 2013). Despite strong evidence that root and shoot trait covariance is an important driver of plant adaptation to drought, few studies have documented how combinations of specific shoot and root traits generate locally adapted ecotypes and the genetic basis of such trait complexes is poorly understood. Genetic crosses and quantitative trait analysis are commonly used tools that can lead to a better understanding of genetic architecture (Michael *et al.*, 2003; Lowry *et al.*, 2014b), and can help to associate genetic variation among and between root and shoot traits with ecotype divergence. Quantitative genetic analyses and the mapping of quantitative trait loci (QTL) permit exploration of the genetic basis of trait correlations and trait divergence (Fishman *et al.*, 2002; Lovell *et al.*, 2015; Milano *et al.*, 2016). Indeed, QTL mapping has been used for identifying the genetic basis of root traits and their function in plant water acquisition (Johnson *et al.*, 2000; Tuberosa *et al.*, 2002; Comas *et al.*, 2013). Importantly, by simultaneously analyzing multiple traits, QTL mapping can infer the loci that drive ecological trait correlations. Functional traits with a high degree of correlation that underlie divergence can result from pleiotropy or genetic linkage through shared developmental genetics (Lande, 1980; Via & Hawthorne, 2005; Lovell *et al.*, 2013) or as a result of correlational selection (Brodie *et al.*, 1995). For example, colocalized QTL for root and shoot traits including root biomass, root volume, shoot biomass and plant height have been identified in a wheat recombinant inbred line population (Iannucci *et al.*, 2017), which may be driven by pleiotropy and / or multiple physically linked genes. Overall, there is growing evidence for substantial genetic variation in root system architecture and root/shoot relationships. However, the loci driving these trait correlations and the degree to which these patterns impact plant productivity are largely unknown.

*Panicum hallii* is a small, self-fertilizing, C_4_ perennial bunch grass native to North America. *P*. *hallii* occurs across a large geographical range with diverse habitats and climates. Average annual precipitation ranges from 127 cm per year on the eastern border of its distribution to 13 cm per year on the west. *P*. *hallii* occurs as two distinct ecotypes (upland and lowland) that are classified as separate subspecies, *P*. *hallii* subsp. *hallii* (hereafter referred to as *hallii)* and *P*. *hallii* subsp. *filipes* (hereafter referred to as *filipes). Hallii* is typically found in upland xeric (habitats with shallow, dry, calcareous and rocky soils in the American southwest and northern Mexico; while *filipes* occurs in lowland mesic areas on clay and silt soils mostly along the Gulf Coast Plain of Texas and Mexico (Gould, 1975; Waller, 1976). The upland ecotype, *hallii*, is smaller in stature and overall size than the lowland ecotype *filipes*: with smaller leaves, fewer tillers, earlier flowering time, less flowers per inflorescence, but larger seed size and seed mass (Waller, 1976; Lowry *et al.*, 2013). This is consistent with its polyploid relative, *Panicum virgatum* (an important biofuel candidate), where upland ecotypes are typically smaller, flower earlier (Lowry *et* al., 2014b) and have lower leaf area (McMillan, 1965) than lowland ecotypes. Previous analyses of shoot traits in a F2 population of *P*. *hallii* (Lowry *et al*., 2014a) demonstrated that a few large-effect loci drove multivariate shoot trait divergence between *hallii* and *filipes*. Here, we investigate the genetic architecture of multidimensional root phenotypic traits and their relationship with shoot phenotypic traits to develop a more complete picture of the adaptive differences between these ecotypes.

In this study, we develop and analyze a recombinant inbred population derived from a cross between the upland and lowland ecotypes of *P*. *hallii*. We employ QTL mapping to identify the genetic architecture of root and shoot trait relationships and their divergence among ecotypes. Specifically, we sought to answer three questions. Are shared QTL involved in genetic correlations between root and shoot traits and biomass partitioning? Do allelic effects of individual QTL underscore root and shoot trait divergence between *hallii* and *filipes*? Will observed QTL and genetic correlations in the RIL population be preserved across two experiments conducted separately under glasshouse and field conditions.

## Materials and Methods

### Development of the RIL mapping population

We developed a population of recombinant inbred lines (RILs) in order to evaluate the genetic basis of divergence between *hallii* and *filipes*. The parents of the RIL mapping population were genotypes selected from populations of the upland and lowland ecotypes of *P*. *hallii*. The upland parent (HAL2-11, hereafter referred to as HAL2) was a one-generation selfed progeny of an individual selected from a glasshouse planting of seed collected from a natural population of *hallii* located at the Lady Bird Johnson Wildflower Center (Austin, TX, USA; 30.16°N, 97.87°W). The lowland parent (FIL2) was selected from a glasshouse planting of seed collected from a natural population of*filipes* located near the coastal city of Corpus Christi, Texas (27.65°N, 97.40°W). FIL2 and HAL2 represent the genome reference genotypes for *filipes* and *hallii* respectively (https://phytozome.jgi.doe.gov/pz/portal.html#!info?alias=Org_Phallii) and are largely homozygous individuals. A cross of these two genotypes, with HAL2 as the maternal parent, yielded an F_1_ hybrid and self-fertilized seed obtained from this individual was used to establish a large F_2_ population (Lowry, 2012). A number of these F_2_ progeny were selected at random and propagated repeatedly via single seed descent until the F_6_ generation. DNA was obtained from leaf tissue of F_6_ seedlings and submitted for whole genome resequencing at the DOE Joint Genome Institute through the Community Science Program. F_7_ seed was subsequently collected from the sequenced F_6_ individuals for this experiment.

SNPs were called from whole genome resequencing of 356 RIL lines on four Illumina 2x150 runs at 12x coverage. Libraries were quality filtered using the fastx toolkit ‘fastq_quality_filter’ program with a quality threshold of 33. Filtered reads were mapped to a soft masked *P*. *hallii* reference genome (FIL-2 V2.0) using bwa mem with the default parameters. Mapped reads were filtered by samtools –Shb with a quality of 20. Bam files were indexed, sorted and duplicates were removed with picard. Reads adjacent to insertions / deletions were masked using GATK RealignerTargetCreator and reads were re-sorted and re-indexed prior to SNP calling. SNPs were called via GATK haplotypeCaller independently for each library, producing a gVCF for each. These were merged and re-genotyped by GATK’s genotypeGVCF and condensed into a 0/1/2 (alternate allele counts) matrix with vcfTools. Genotype data from 335 RIL lines were included in the output genotype matrix. The resultant matrix was processed in R. SNPs with >10% and <80% homozygotes and <5% NA and <20% heterozygotes were retained.

We applied a 3-step sliding window approach for marker calling: 1) The genome was broken into 200 marker windows (overlapping by 100 markers) and the proportion of each genotype was calculated. 2) Training data was constructed, retaining the 100 strongest heterozygous sites and a random sampling of 100 of the sites with > the mean proportion of each homozygote; 3) A random forest machine learning model was fit to the training data (the R caret package) and used to predict the genotypes of all sliding window intervals resulting in a 3361 marker matrix.

### Genetic map construction

To build the genetic map, we culled the genotype matrix such that no two markers could have a pairwise recombination fraction <0.005. This culling procedure minimized the amount of segregation distortion and missing data within any 0.5 cM window. Linkage groups were formed from the resulting 1278 marker matrix. Markers were ordered within linkage groups using a travelling salesperson problem solver as implemented through the concorde program and parsed through the TSPMap function tspOrder (Monroe *et al.*, 2017). We then fine-tuned the resulting genetic map first by culling the genotype matrix to a 711-marker grid where no markers resided <1cM from an adjacent marker, then looking at improving the fine-order of markers using the ripple algorithm. Finally, chromosomes were named and oriented to maximize the similarity with the physical position of markers in the FIL2 genome annotation (phytozome.net).

### Morphological shoot and root phenotyping under glasshouse conditions

Seeds of 174 F_7_ RILs and the two parental genotypes were scarified with sandpaper and placed on wet sand in round petri dishes on September 5, 2016 and allowed to germinate in a glasshouse located at the University of Texas at Austin, Brackenridge Field Lab (12-h days at 500 μE m ^−2^ s ^−1^, 28°C; 12-h nights at 24°C). On the 7^th^ day after sowing, seedlings were transferred to 6 cm x 30 cm cone-tainers (Stuewe and Sons, Tangent, OR). Cone-tainers were lined with 1 mil plastic liners (perforated at the bottom for drainage) to facilitate separation of the plant from the container during harvest. Cone-tainers were filled with Field and Fairway Profile (The Turf Trade, NJ, USA) media. Plants were then assigned to a completely randomized block design within three blocks on a single glasshouse bench. Plants were bottom watered every three days with Grow liquid nutrient solution (DynaGro, Richmond, CA) to promote seedling growth. Plants were harvested within three days of a common developmental stage defined as when a fully expanded flag leaf with a visible ligule was observable on any tiller with an emerging panicle. The plant in its plastic bag was pulled from the pot gently to prevent damage to the root system. Then the bag was cut open and the profile substrate was gently removed by shaking the plant on wire mesh followed by light washing of the root system in a bucket of tap water. Shoot material was separated from root material. The tiller height (from base of the plant to the node of the flag leaf on the tiller with the emergent panicle), leaf length and area of the flag leaf of the main tiller were measured and tiller number was counted at the time of harvest. Total root number was counted and then the root system was spread out in a clear acrylic water filled tray and scanned at a 600 dpi resolution using an EPSON Scanner (Model 12000XL, Epson America, Inc., San Jose, CA, USA) calibrated for use with WinRhizo Pro 2015 root image analysis software (Regent Instruments Inc., Canada). Leaf, shoot and root tissue was collected separately, dried for 96 hours in an oven at 55°C, and weighed to obtain biomass. Specific leaf area (SLA) was calculated as fresh leaf area divided by dry mass of the leaf (cm^2^ g^-1^).

Root trait data was obtained from scans using WinRhizo Pro 2015 software and included total root length (cm), total root volume (cm^3^), and average root diameter (mm). The Lagarde’s local threshold parameter in the analysis software was enabled to ensure detection of thin and pale roots and the diameter class size was set to 0.25 mm to ensure accurate calculation of average root diameter. Specific root length (SRL; total root length / root biomass (cm g^-1^)), root tissue density (root biomass / total root volume (g cm^-3^)), root length density (total root length / soil volume (cm cm^-3^)) and root mass ratio (RMR, root biomass/total biomass) were calculated for each plant.

### Data and QTL analysis

Data analyses centered on fitting linear mixed models and considered RIL genotype as a fixed effect (proc mixed, SAS) for the measured shoot and root phenotypic traits. Block was also included as a fixed effect covariate when it had a significant impact on measured traits (emergence day, specific root length and root diameter). The SAS procedure PROC CORR was used to calculate genetic correlation coefficients of traits based on RIL line means. Trait heritability was calculated using h2boot software with one-way ANOVA among clonal lines with 1000 bootstrap runs (Phillips & Arnold, 1999). Trait divergence between parental lines was tested with a t-test in SAS.

The majority of the measured traits were continuously distributed with relatively strong multivariate structure based on pairwise correlational analyses. As such, we also used genetic principal component analysis (PCA) to obtain a multidimensional overview of shoot and root trait variation and integration. PCA analysis was performed on the trait means of each line for the following phenotypic variables: emergence day, tiller number, root number, root biomass, shoot biomass, root diameter, root tissue density, specific root length, specific leaf area, tiller height, leaf length, root volume and total root length. PCA was completed using SAS with the proc princomp function. The first three principal components that together explained 75% of total variation were retained for QTL analysis.

QTL mapping was completed in R using the R/qtl package (Broman & Sen, 2009) on the RIL breeding values as described above (Table S2). When quantitative trait data distributions were not normally distributed, data was log (emergence day, tiller number) or square root (shoot biomass) transformed. Two functions were used to determine the position of QTL and to conduct the calculation of estimates for additive effects and effects of epistasis (an additive-by-additive interaction between quantitative trait loci, script: https://github.com/AlbinaKh/P.hallii_RIL_RootShoot_QTLmapping). The scantwo function with 1000 permutations was used to calculate penalties for main effect and interactions for each phenotypic trait, and the stepwise QTL function was used to conduct a forward-backward search and account for epistasis with a maximum of 6 QTL (at least two QTL peaks in addition to those detected with the scanone function) that optimized the penalized LOD score criterion. Threshold values for type 1 error rates were set at alpha = 0.05 for all traits based on permutation. 1.5 LOD drop interval of QTL was calculated using the qtlStats function. QTL analysis was performed on the first three principal components following the above procedure.

### Confirming root and shoot biomass QTL in a field study

An important question in root biology is the degree to which genetic effects discovered in controlled conditions (i.e. glasshouse, growth chamber, or artificial media experiments) relate to real world variation in root system architecture in the field (Champoux *et al.*, 1995; Ochoa *et al.*, 2006; Lovell *et al.*, 2016). To further confirm and evaluate major QTL detected in our glasshouse study, we conducted a follow up experiment on a focal QTL during the 2016 growing season. Ten RILs homozygous at the shared QTL region for root and shoot biomass were selected for this experiment (5 with *filipes* alleles and 5 with *hallii* alleles). Seed of selected lines were germinated and established in the glasshouse using the procedure outlined above for the RIL planting and subsequently transplanted into the field at the age of one month. Eight biological replicates of each line and eight replicates of the parental genotypes were planted on May 10, 2016 under both restrictive and well-watered irrigation treatments (10 RIL lines x 2 parents x 8 biological replicates x 2 irrigation levels = 192 plants). All plants were well-watered for one week after transplant for establishment.

The field experiment was conducted at a site located within the Brackenridge Field Laboratory property of the University of Texas in Austin, TX, USA (N 30.2845, W 97.7809). The site elevation is 133 m above sea level and soils are Yazoo sandy loam greater than 1.2 m deep. The mean maximum temperature (August) is ~35.0 °C and the mean minimum temperature (January) is ~ 3.0 °C. This experiment was co-planted in available space within an existing *P*. *hallii* experiment to take advantage of established irrigation infrastructure. The site contains 32 differentially irrigated ‘beds’ which are separated underground by 1.2-meter-deep plastic sheeting (Regal Plastics, Austin, TX, USA) to prevent the spread of applied irrigation water. Irrigation was applied by dripline (0.9 GPH, 12” emitter spacing, Rain Bird, Azusa, CA). The treatment period occurred from June through August with the restrictive treatment receiving 4.5 fold less irrigation in both number of irrigation events and total amount of water applied. Plants were harvested towards the end of the summer growing season in August over a three-day period. To account for differences in size of the plants, an equal volume of the soil under each plant was harvested using a ‘shovelomics’ device that regulated shovel angle and depth while extracting plants from the field soil. Plants with roots attached were rinsed clean of soil over a metal screen and allowed to air dry. Shoot biomass was then separated from root biomass, and both biomass types were dried at 55°C for 4 days before weighing. Trait values more extreme than 1.5x the interquartile range were removed as outliers prior to analysis. For statistical analysis, we used linear mixed models with proc mixed in SAS. The main effect for the model was genotype of the QTL (*filipes* or *hallii* alleles at the marker position), treatment and genotype by treatment interaction. RIL line was used as a random effect to control for background genetic variance.

## Results

### Heritable shoot and root trait differences between upland and lowland ecotypes

The RIL parents representing upland and lowland ecotypes of *Panicum hallii* (HAL2 and FIL2) had significantly different shoot trait mean values (Table 1) and also displayed corresponding differences between root traits. The upland genotype, HAL2, had 2.3-fold faster first panicle emergence (*t* values at 5 dfs and *P* values; *t*=2.87, *P*=0.035), 3.3-fold less shoot biomass (*t*=4.39, *P*=0.007) and 2.8-fold less root biomass (*t*= 3.08, *P*=0.028), 1.8-fold shorter plant height (*t*= 3.43, *P*=0.018), 2.2-fold shorter leaf length (*t*=6.3, *P*=0.001), 2-fold shorter total root length (*t*=3.29, *P*=0.022), 2.5-fold lower total root volume (*t*=3.41, *P*=0.02), and 1.3-fold increased specific root length (*t*=-2.5, *P*=0.05) relative to the lowland genotype FIL2 (Table 1).

**Table 1.** Means, one standard error (SE) and broad-sense heritability (*H^2^)* of root and shoot traits for the *Panicum hallii* RIL population and its parental genotypes.

| Phenotypic Trait | FIL2 | HAL2 | *t* | *P*-value | RIL mean | RIL range | $H^2 \pm SE$ |
| --- | --- | --- | --- | --- | --- | --- | --- |
| Panicle Emergence (day) | 9.25±1.19 | 4.00±1.38 | 2.87 | **0.035** | 7.01±1.74 | 1.00 – 18.33 | 0.51±0.05 |
| Shoot Biomass (g) | 4.74±0.49 | 1.41±0.57 | 4.39 | **0.007** | 1.65±0.33 | 0.29– 4.74 | 0.59±0.05 |
| Tiller Number | 6.25±0.48 | 5.00±0.56 | 1.68 | 0.150 | 6.00±0.83 | 3.00 – 14.50 | 0.50±0.05 |
| SLA | 325.62±18.15 | 382.77±20.96 | -2.06 | 0.094 | 381.58±33.17 | 264.67 – 499.36 | 0.18±0.08 |
| Plant Height (cm) | 21.18±1.82 | 11.63±2.11 | 3.43 | **0.018** | 12.57±1.56 | 4.30 – 23.65 | 0.63±0.04 |
| Leaf Length (cm) | 30.77±1.72 | 14.23±1.98 | 6.30 | **0.001** | 15.66±1.46 | 4.75– 24.27 | 0.66±0.04 |
| Root Biomass (g) | 1.38±0.18 | 0.51±0.21 | 3.08 | **0.028** | 0.54±0.10 | 0.12 – 1.60 | 0.58±0.06 |
| Root Number | 14.00±0.97 | 8.33±1.11 | 3.84 | **0.012** | 8.87±1.39 | 2.50 – 15.00 | 0.38±0.05 |
| SRL (cm g<sup>-1</sup>) | 10.14±0.85 | 13.37±0.98 | -2.50 | 0.055 | 12.27±1.11 | 6.12 – 17.95 | 0.43±0.06 |
| RTD (g cm<sup>-3</sup>) | 0.06±0.01 | 0.05±0.01 | 1.31 | 0.247 | 0.05±0.01 | 0.03 – 0.08 | 0.39±0.07 |
| Root Diameter (mm) | 0.46±0.01 | 0.44±0.02 | 1.27 | 0.259 | 0.45±0.01 | 0.37 – 0.55 | 0.37±0.05 |
| Root Volume (cm<sup>3</sup>) | 2.43±0.28 | 0.98±0.32 | 3.41 | **0.019** | 1.00±0.17 | 0.26 – 2.90 | 0.56±0.05 |
| Root Length (m) | 1.37±0.14 | 0.67±0.16 | 3.29 | **0.022** | 0.65±0.11 | 0.12 – 1.64 | 0.59±0.04 |
| RMR | 0.22±0.01 | 0.27±0.01 | -3.44 | **0.018** | 0.25±0.02 | 0.16 – 0.39 | 0.34±0.09 |

FIL2, *P. hallii* subsp. *filipes*; HAL2, *P. hallii* subsp. *hallii*. Values are mean ± 1SE. *t*, t-statistics at 5 degrees of freedom in test for divergence
between parental lines. Statistically significant values are indicated in bold text.

We estimated broad-sense trait heritability (*H^2^*) as the proportion of observed phenotypic variance due to genetic differences among RILs in the population. In the RIL population, all measured traits were heritable, with *H^2^* ranging from 18% to 66% for shoot traits (bootstrap based significance, in all cases *P*<0.001) and from 34% to 60% for root traits (bootstrap bases significance, in all cases *P*<0.001). The most heritable traits were leaf length (66%), plant height (64%), shoot biomass (60%), root length (60%) and root biomass (58%; Table 1). Transgressive segregation, where the range of recombinant phenotypes extends beyond the range of parental values (Rieseberg *et al.*, 1999), was found among the majority of traits except shoot biomass, plant height, leaf length, root biomass and root number. In the parental lines, FIL2 had trait values that were the highest or close to the highest of population wide values, while HAL2 values were generally in the middle of the population trait distribution (Table 1).

Many shoot and root phenotypic traits also showed remarkably strong genetic correlations in the RIL population (Table 2). For example, shoot and root biomass (r=0.92, *P*<0.0001), tiller and root number (r=0.67, *P*<0.001), shoot biomass and root volume (r=0.91, *P*<0.0001), and shoot biomass and total root length (r=0.90, *P*<0.001). We performed principal component analysis (PCA) to characterize the multivariate structure of our data. The first three PCA axes explained 75% of the overall trait variance. Principal component one (PC1; 45.5% variance explained) was composed of general plant size traits. Principal component two (PC2; 16.5%) was mainly composed of root resource acquisition traits. Principal component three (PC3; 12.6%) was composed of carbon acquisition and allocation traits (Table S1; Fig. 1a, b, c).

**Table 2.** Pearson Correlation Coefficients for genetic correlations in the *Panicum hallii* RIL population.

| Trait | ED | TN | RTN | SHMASS | RTMASS | SRL | RTD | HEIGHT | LFLG | RMR | SLA | RTDM | RTLG |
| --- | --- | --- | --- | --- | --- | --- | --- | --- | --- | --- | --- | --- | --- |
| TN | 0.116 |  |  |  |  |  |  |  |  |  |  |  |  |
| RTN | 0.039 | **0.669** |  |  |  |  |  |  |  |  |  |  |  |
| SHMASS | **0.215** | **0.545** | **0.758** |  |  |  |  |  |  |  |  |  |  |
| RTMASS | 0.132 | **0.615** | **0.789** | **0.921** |  |  |  |  |  |  |  |  |  |
| SRL | **-0.192** | **-0.115** | -0.05 | -0.021 | **-0.15** |  |  |  |  |  |  |  |  |
| RTD | **0.195** | **0.195** | **0.191** | **0.281** | **0.336** | **-0.544** |  |  |  |  |  |  |  |
| HEIGHT | 0.119 | **0.285** | **0.598** | **0.824** | **0.719** | 0.118 | 0.136 |  |  |  |  |  |  |
| LFLG | 0.015 | **0.2** | **0.573** | **0.759** | **0.678** | 0.135 | 0.109 | **0.769** |  |  |  |  |  |
| RMR | **-0.281** | -0.005 | -0.121 | **-0.424** | -0.085 | **-0.267** | 0.029 | **-0.495** | **-0.399** |  |  |  |  |
| SLA | **-0.331** | 0.079 | 0.001 | -0.223 | -0.095 | **0.259** | **-0.288** | -0.114 | -0.047 | **0.388** |  |  |  |
| RTDM | 0.125 | -0.071 | **-0.183** | **-0.260** | **-0.163** | **-0.696** | -0.115 | **-0.317** | **-0.338** | **0.290** | -0.135 |  |  |
| RTLG | 0.057 | **0.566** | **0.772** | **0.905** | **0.925** | **0.185** | **0.198** | **0.762** | **0.734** | **-0.196** | 0.003 | **-0.450** |  |
| RTVOL | 0.120 | **0.614** | **0.802** | **0.911** | **0.975** | -0.059 | **0.150** | **0.722** | **0.670** | -0.107 | -0.045 | -0.134 | **0.934** |

ED, panicle emergence; TN, tiller number; RTN, root number; SHMASS, shoot biomass; RTMASS, root biomass; SRL, specific root length; RTD, root tissue density; HEIGHT, plant height; LFLG, leaf length; RMR, root mass ratio; SLA, specific leaf area; RTDM, root diameter; RTLG, root length. Significant correlations are indicated in bold text.

**Fig. 1.**
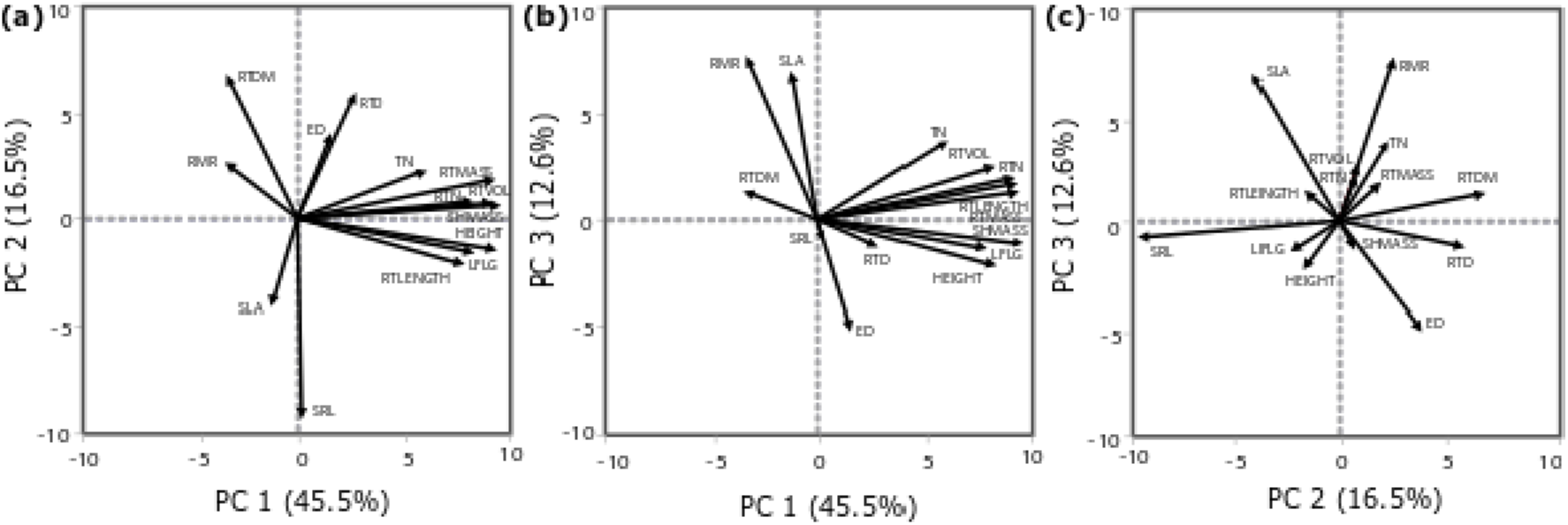
Principle component analysis of shoot and root traits for the *Panicum hallii* RIL population. Traits: PC, principal component; RMR, root mass ratio; SLA, specific leaf area; SRL, specific root length; RTLRNGTH, root length; LFLG, leaf length; HEIGHT, plant height; SHMASS, shoot biomass; RTMASS, root biomass; RTVOL, root volume; RTN, root number; TN, tiller number; RTD, root tissue density; ED, emergence day; RTDM, root diameter.

### QTL underscore root and shoot trait divergence between hallii and filipes

Given high *H^2^* values, it is not surprising that QTL were detected for all measured traits. A total of 31 QTL were identified for 13 phenotypic traits: two for one phenological trait, 14 QTL for five shoot traits and 15 QTL for seven root traits (Table 3, Fig. 2). QTL for all traits showed additive effects in the direction of parental divergence, except for one of three QTL for tiller number, one of four QTL for root diameter, and one of three QTL for SRL. *Filipes* alleles increased trait values associated with phenology and plant size, including: emergence day, root number, root tissue density, root biomass, shoot biomass, tiller height, leaf length and root volume. *Hallii* alleles increased trait values associated with resource acquisition and allocation, including: specific root length, root mass ratio, and specific leaf area. The main effects of each QTL explained from 5.25% to 15.4% of phenotype variation for shoot traits, and from 5.9% to 18.6% for root traits (Table 3). Of these 31 QTL, eight QTL occupied unique positions throughout genome: RTD on chr1, leaf length on chr2, tiller number on chr3 and chr8, root number on chr3, SLA on chr5, tiller height and root diameter on chr8. Three of these single QTL were also confirmed by principle component QTL (Table 4, Fig. 2). The confidence intervals of all other QTL are shared with at least one other QTL.

**Table 3.** Main and epistatic effects of QTL for the *Panicum hallii* RIL population.

| Phenotype | Chr | Peak<br>(cM) | 1.5 Lod<br>Interval | LOD | %<br>var | Effect | SE | Donor<br>of<br>Positive<br>allele | QTL<br>Cluster<br>(CL) |
| --- | --- | --- | --- | --- | --- | --- | --- | --- | --- |
| Panicle<br>Emergence<br>(day) | 5 | 52.1 | 40-59 | 4.59 | 9.85 | -0.044 | 0.009 | *filipes* | CL5.1 |
|  | 7 | 80.0 | 31-83 | 4.31 | 9.2 | -0.039 | 0.008 | *filipes* | CL7.2 |
| Shoot Biomass<br>(g) | 5 | 58.6 | 56-60 | 7.43 | 14.8 | -0.044 | 0.007 | *filipes* | CL5.1 |
|  | 5 | 136.0 | 128-142 | 5.08 | 9.82 | -0.031 | 0.007 | *filipes* | CL5.3 |
|  | 9 | 66.1 | 60-71 | 4.78 | 9.19 | -0.027 | 0.005 | *filipes* | CL9.1 |
|  | Epi5:5 |  |  | 2.86 | 5.36 | 0.027 | 0.007 |  |  |
| Tiller Number<br>(count) | 3 | 40.5 | 38-48 | 7.23 | 14.74 | -0.054 | 0.009 | *filipes* |  |
|  | 5 | 137.0 | 128-142 | 3.47 | 6.73 | -0.037 | 0.009 | *filipes* | CL5.3 |
|  | 7 | 73.6 | 46-81 | 4.84 | 9.56 | 0.039 | 0.008 | *hallii* | CL7.2 |
| SLA<br>(cm<sup>2</sup>g<sup>-1</sup>) | 5 | 13.3 | 0-26 | 3.15 | 5.25 | 9.772 | 2.543 | *hallii* |  |
|  | 7 | 66.0 | 60-74 | 8.56 | 15.37 | 16.394 | 2.494 | *hallii* | CL7.2 |
|  | 8 | 19.8 | 16-23 | 8.33 | 14.90 | 16.077 | 2.484 | *hallii* |  |
| Tiller Height<br>(cm) | 5 | 76.0 | 74-77 | 6.16 | 13.34 | -1.765 | 0.320 | *filipes* | CL5.2 |
|  | 6 | 83.9 | 69-88 | 3.82 | 8.05 | -1.096 | 0.256 | *filipes* |  |
| Leaf Length<br>(cm) | 2 | 89.7 | 76-96 | 4.28 | 8.56 | -1.19 | 0.264 | *filipes* |  |
|  | 7 | 43.6 | 35-64 | 4.39 | 8.80 | -1.293 | 0.283 | *filipes* | CL7.1 |
|  | 9 | 63.4 | 59-75 | 3.41 | 6.76 | -0.985 | 0.246 | *filipes* | CL9.1 |
| Root Biomass<br>(g) | 5 | 58.6 | 56-60 | 8.81 | 18.61 | -0.012 | 0.002 | *filipes* | CL5.1 |
|  | 5 | 136.0 | 135-142 | 8 | 16.71 | -0.010 | 0.002 | *filipes* | CL5.3 |
|  | Epi5:5 |  |  | 4.61 | 9.21 | 0.008 | 0.002 |  |  |
| Root Number<br>(count) | 3 | 88.0 | 69-104 | 6.18 | 13.9 | -1.08 | 0.199 | *filipes* |  |
|  | 5 | 125.7 | 125-130 | 5.36 | 11.94 | -0.81 | 0.196 | *filipes* | CL5.3 |
|  | Epi3:5 |  |  | 2.79 | 5.99 | 0.73 | 0.202 |  |  |
| SRL (cm g<sup>-1</sup>) | 1 | 91.5 | 82-94 | 5.3 | 11.02 | 0.66 | 0.131 | *hallii* | CL1.1 |
|  | 3 | 18.8 | 17-36 | 5.16 | 10.7 | 0.78 | 0.156 | *hallii* | CL3.1 |
|  | 7 | 44.7 | 34-49 | 3.16 | 6.4 | -0.55 | 0.145 | *filipes* | CL7.1 |
| RTD (g cm<sup>-3</sup>) | 1 | 6.3 | 0-20 | 3.15 | 7.9 | -0.001 | 0.0004 | *filipes* |  |
| Root Diameter<br>(mm) | 1 | 86.0 | 82-94 | 4.73 | 8.68 | -0.009 | 0.002 | *filipes* | CL1.1 |
|  | 3 | 34.2 | 30-36 | 5.36 | 9.91 | -0.011 | 0.002 | *filipes* | CL3.1 |
|  | 5 | 71.9 | 66-75 | 3.78 | 6.84 | 0.010 | 0.002 | *hallii* | CL5.2 |
|  | 8 | 47.9 | 43-52 | 4.65 | 8.50 | -0.009 | 0.002 | *filipes* |  |
| Root Volume<br>(cm<sup>3</sup>) | 5 | 58.6 | 56-63 | 3.96 | 8.85 | -0.134 | 0.030 | *filipes* | CL5.1 |
|  | 5 | 117.2 | 109-142 | 3.07 | 6.77 | -0.119 | 0.032 | *filipes* | CL5.3 |
| Root Length | 5 | 58.6 | 44-138 | 3.12 | 7.85 | -0.82 | 21.29 | *filipes* | CL5.1,2,3 |
| RMR (ratio) | 7 | 67.0 | 62-74 | 6.36 | 15.34 | 0.0137 | 0.002 | *hallii* | CL7.2 |

Chr, chromosome; Peak, cM (centimorgan) position of the QTL peak; LOD, logarithm of odds; % var, present of variance; SE, one standard error; SLA, specific leaf area; SRL, specific root length; RTD, root tissue density; RMR, root mass ratio; Epi, epistasis.

**Fig. 2.**
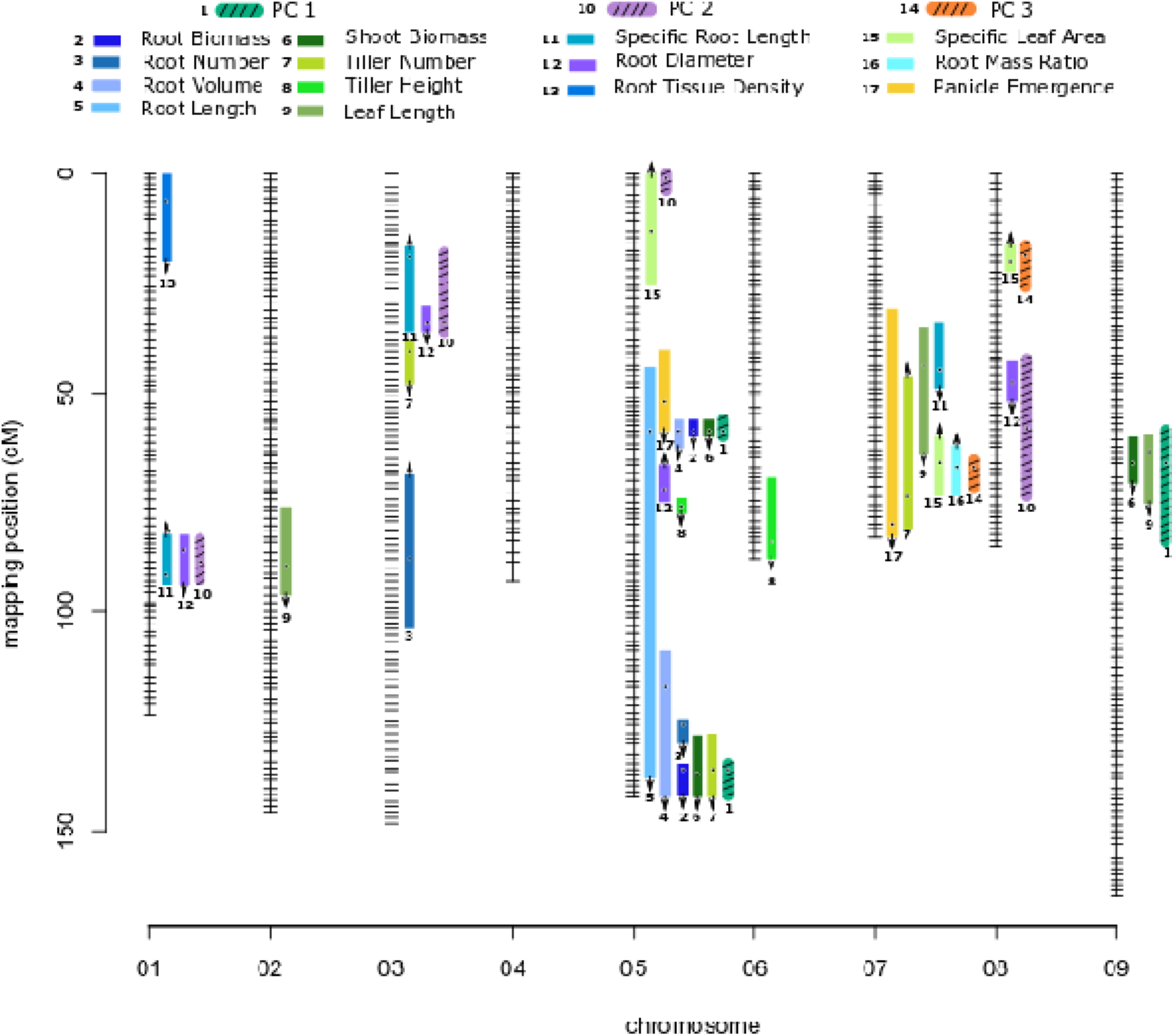
Genetic map of the *Panicum hallii* RIL population with location of trait QTL. Colored bars indicate 1.5-LOD drop confidence intervals. Location of dots within the bars is the location of QTL peaks. Arrow is the direction of additive effect, with up or down arrow indicating the HAL2 allele increases or decreases the trait value.

### Trait-specific QTL cluster into genomic ‘hotspots’

Divergence of correlated traits in natural populations may be driven by pleiotropic genes or linked genes with correlated effects. We identified three major and five minor clusters of root and shoot trait QTL over five different chromosomes (Table 3, Fig. 1). Here we identify clusters (CL) by chromosome number and number from the 0 cM position for each chromosome. As expected, we found that positions of QTL for principle components were highly indicative of the locations of QTL clusters for the traits included in their particular PC axis (Table 4, Fig. 2).

**Table 4.** Main and epistatic effects of the first three principal component QTL for the *Panicum hallii* RIL population.

| Principal Component | Chr | Peak (cM) | 1.5 Lod Interval | LOD | % var | Effect | SE | Donor of Positive allele | QTL Cluster (CL) |
| --- | --- | --- | --- | --- | --- | --- | --- | --- | --- |
| PC1 | 5 | 58.6 | 56-60 | 7.19 | 14.42 | -1.215 | 0.209 | *filipes* | CL5.1 |
|  | 5 | 136.0 | 135-142 | 6.34 | 12.67 | -1.049 | 0.206 | *filipes* | CL5.3 |
|  | 9 | 66.1 | 58-84 | 3.51 | 6.7 | -0.664 | 0.164 | *filipes* | CL9.1 |
|  | Epi5:5 |  |  | 3.14 | 6.0 | 0.812 | 0.212 |  |  |
| PC2 | 1 | 88.7 | 83-93 | 4.51 | 8.26 | -0.458 | 0.099 | *filipes* | CL1.1 |
|  | 3 | 34.2 | 18-36 | 4.97 | 9.14 | -0.533 | 0.109 | *filipes* | CL3.1 |
|  | 5 | 1.1 | 0-4 | 4.13 | 7.52 | 0.457 | 0.103 | *filipes* |  |
|  | 8 | 58.0 | 42-74 | 3.45 | 6.24 | 0.392 | 0.097 | *filipes* |  |
| PC3 | 7 | 67.0 | 65-72 | 12.14 | 25.05 | 0.676 | 0.084 | *hallii* | CL7.2 |
|  | 8 | 18.5 | 16-26 | 3.59 | 6.60 | 0.354 | 0.085 | *hallii* | CL8.1 |

Chr, chromosome; Peak, cM (centimorgan) position of the QTL peak; LOD, logarithm of odds; % var, present of variance explained; SE, one standard error; PC1, principal component 1; PC2, principal component 2; PC3, principal component 3; Epi, epistasis.

PC1 QTL associated with three clusters of QTL for plant size traits. CL9.1 contains shoot biomass and leaf length. CL5.1 contains root biomass, shoot biomass, root volume, and panicle emergence and CL5.3 contains root biomass, shoot biomass, root volume, tiller number and root number. Both chr5 clusters overlap the large confidence interval of the QTL for total root length. To test if there are multiple undetected QTL for total root length that could explain the large confidence interval, we lowered the LOD threshold and found two separate QTL associated with each of the two primary chr5 clusters (data not shown). A separate QTL pair of tiller height and root diameter not indicated by PC1 lies between these two large clusters. PC2 QTL associated with of two clusters of QTL for root resource acquisition traits. CL1.1 and 3.1 both contain SRL and root diameter. PC3 associated with a single cluster (CL7.2) related to carbon allocation traits. CL7.2 contains panicle emergence day, leaf length, number of tillers, RMR and SLA. Near this PC3 associated QTL is a minor cluster (CL7.1) of leaf length and SRL (Fig. 2). Four pairwise epistatic interactions, where the effect of one QTL depends on the allelic state of an unlinked QTL, were detected (Table 3, 4; Fig. 3 a-d). Three QTL from cluster CL5.3 (shoot biomass, root biomass and PC1) interacted with other QTL for these traits located in CL5.2; and the root number QTL from CL5.3 interacted with the root number QTL on chr3. Individuals that possess the *hallii* allele for these QTL at CL5.3 mask the positive effects of their interactive QTL.

**Fig. 3.**
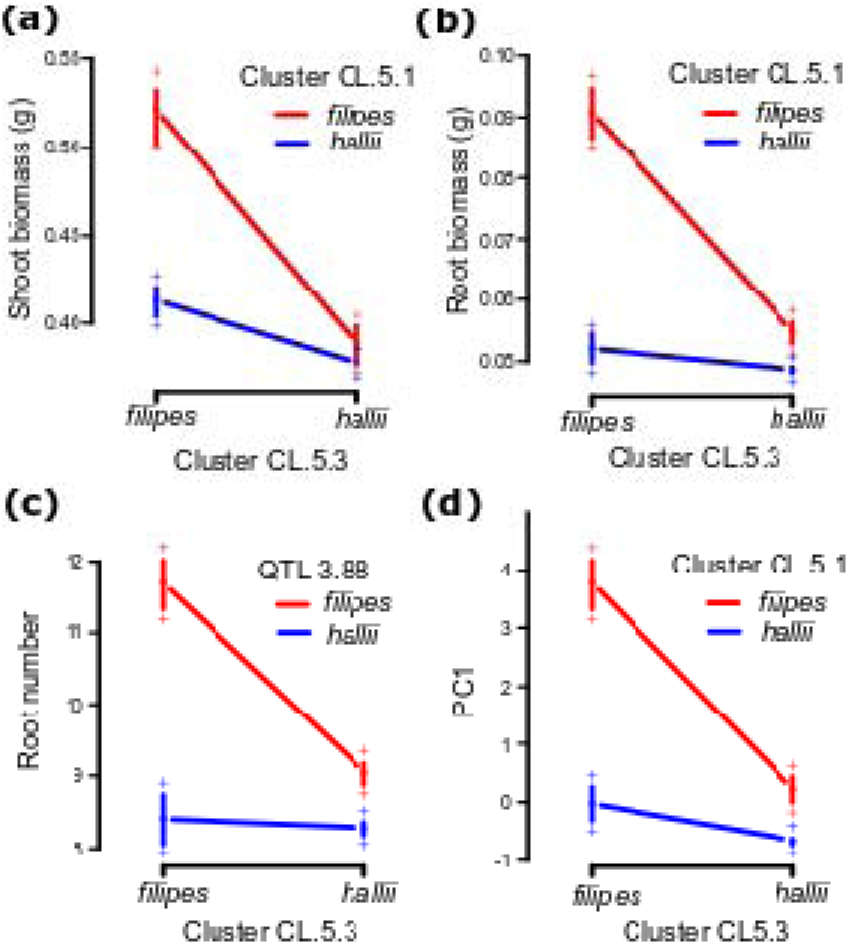
Pairwise epistatic QTL in the *Panicum hallii* RIL population. Plotted points indicate two-locus genotype means ± 1SE for the two loci containing root biomass (a), shoot biomass (b), root number (c) and PC1 (d).

**Fig. 4.**
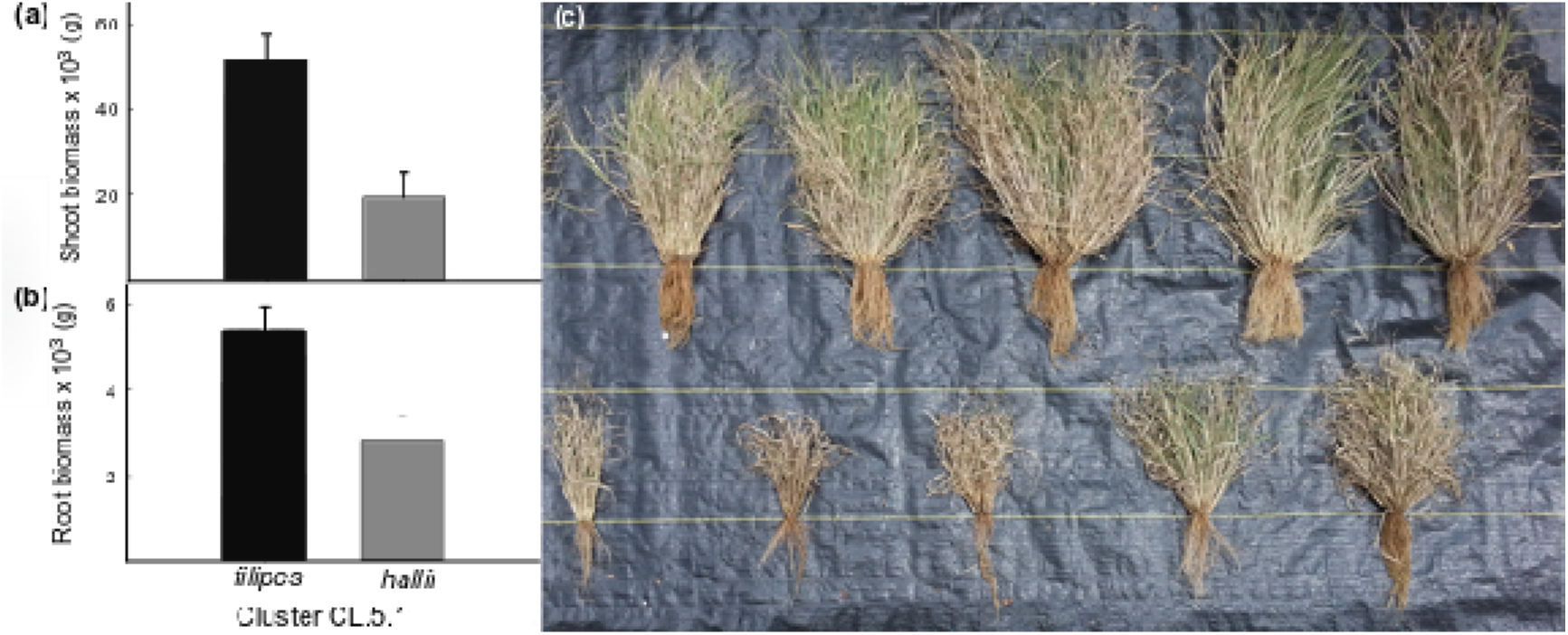
Mean ± 1SE of shoot biomass (a) and root biomass (b) for field grown *Panicum hallii* RIL lines homozygous for either *filipes* or *hallii* parental alleles at shoot and root biomass QTL located in cluster CL5.2. Picture of field grown RIL lines homozygous at CL5.2 for *filipes* allele (top row) and *hallii* allele (bottom row) (c).

### A Major Pleotropic Effect QTL is Confirmed in the Field

To confirm the effects of QTL observed in a controlled glasshouse study, we phenotyped two sets of RIL lines homozygous for different parental alleles at the loci for shoot and root biomass (CL5.2) in a field experiment. While the magnitude of increased biomass for lines with *filipes* alleles at the selected QTL observed in the field is 24% less for the root biomass and 11% less for the shoot biomass relative to the glasshouse, the effects are significant and in the same direction as those observed in the glasshouse. Field grown lines with *filipes* parental alleles produced 1.9-fold more root biomass (*P*=0.0024) and 2.7-fold more shoot biomass (*P*=0.0002) relative to field grown lines with *hallii* parental alleles (Fig. 4). In addition, the HAL2 parental line showed a 1.8-fold increase trend in RMR (*P*=0.09) over the FIL2 parental line under field conditions compared to the 1.2-fold difference observed in the glasshouse (*P*=0.018). There were no significant differences between the irrigation treatments or the interaction of treatment by genotype for RIL lines or the parental genotypes. However, root biomass showed a 1.2-fold increase trend under the dry treatment relative to the wet treatment (*P*=0.08).

## Discussion

Previous tests of upland and lowland ecotype divergence have focused primarily on shoot traits and their relationship to plant water status and drought strategies (Maherali *et al.*, 2008; Latta 2009; Olsen *et al.*, 2013; Lowry, 2014a). Root traits can also be involved in adaptive differentiation to abiotic stresses by their direct effects on water acquisition, and through correlation, tradeoffs or constraints with shoot traits (Hammer *et al.*, 2009; Mace *et* al., 2012). Understanding the genetic control of root and shoot trait integration will aid in developing a more complete picture of the process of ecotype divergence. In this study, to examine both above and below ground trait divergence, we assembled a high quality genetic linkage map by resequencing a recombinant inbred population derived from a cross between upland and lowland ecotypes of *Panicum hallii* and conducted a QTL analysis. We mapped at least one QTL for all measured shoot and root traits. QTL for the majority of ecotype differentiating root and shoot size traits were colocalized into several genomic ‘hotspots’ while QTL for differentiating resource acquisition traits mapped to largely independent genomic regions. We characterized allelic effects of individual QTL for traits involved in ecotype differentiation and found that they underscored trait divergence between the parental populations. A subsequent field study conducted on fully mature individuals at the end of the growing season confirmed the co-localized QTL relationships for shoot and root biomass that were observed in the glasshouse study of plants at first maturity.

### Genetic architecture of shoot and root traits underlying ecotype divergence

Ecotypes are often differentiated by suites of correlated root and shoot traits that may share common genetic and developmental architectures as a result of adaptive differentiation. One of our major findings was several genomic ‘hotspots’ of colocalized QTL for multiple shoot and root traits. This is consistent with the previous study of a *P*. *hallii* F_2_ population covering a suite of ecotype differentiating shoot trait QTL which clustered on chr5 (Lowry et al. 2014a). In addition to confirming this important locus, we discovered additional root traits linked to this region and additional regions of clustered loci for root and shoot traits. This pattern of colocalized QTL that control traits such as root biomass, shoot biomass, among others, has also been shown in a RIL populations of wheat and sorghum (Iannucci *et al.*, 2017; Mace *et al.*, 2012). These findings indicate that specific loci can shape both shoot and root morphological traits, through tight linkage of several genes controlling individual traits or a single pleiotropic gene that controls several traits. All QTL clusters detected in this RIL population were indicated by various QTL for principle components.

PC1 indicated three QTL clusters of multiple size related root and shoot traits (shoot biomass, root biomass, root volume, and other). We found that the *hallii* allele had additive effects in the direction of ecotype divergence and contributed to smaller root and shoot phenotypes in every case compared to the *filipes* allele. This finding is consistent with the global pattern observed in angiosperm plants whose shoot and root biomass are positively correlated (Enquist & Niklas, 2002) and with other studies on perennial grasses where total biomass is decreased under water limited conditions (Baruch, 1994; Weißhuhn *et al.*, 2011; Tozer *et al.*, 2017).

In addition to differences in absolute size, there are expected differences in carbon acquisition and allocation between upland and lowland ecotypes. PC3 indicated one cluster of carbon allocation and phenology related traits (SLA, RMR, tiller number and panicle emergence). Plants with *hallii* alleles had greater SLA, RMR, tiller number, and faster panicle emergence. Thinner leaves (high SLA) are less carbon costly to produce and associated with increased photosynthetic capacity (Reich *et* al., 1997; Cornelissen *et al.*, 2003). Increased RMR helps to maintain plant water status and productivity under drought (Comas *et al.*, 2013). Faster flowering time along with greater tiller number allows for rapid production of seeds when resources are available for short time periods. These factors combined may indicate that *hallii* employs a fast acquisitive strategy for drought escape; acquiring nutrients rapidly and flowering quickly to enter a dormant state before periods of summer drought. Acquisitive shoot and root strategies have been associated with fast growth strategies and summer dormancy in other perennial grasses (Balachowski *et al.*, 2016). This contrasts with the lower SLA, and RMR of the lowland *filipes*, which may employ a slow strategy of thicker longer lasting leaves, larger more persistent roots, and abundant above ground foliage. This common genetic control of ecotype differentiating traits involving shoot and root organs suggests that these factors evolved in tandem.

SLA and SRL are both important plant traits that are linked to resource acquisition (Reich, 2014; Cheng *et al.*, 2016) and associated with fast growth (Reich, 2014, Pérez-Harguindeguyb *et al.*, 2016). SRL is typically thought of as the below ground analog of SLA (Eissenstat *et al.*, 2000; Reich, 2014), where an acquisitive root strategy (high SRL) can be aided by an acquisitive leaf strategy (high SLA, Perez-Ramos *et al.*, 2013). In some cases, these traits are found to be positively correlated (Withington *et al.*, 2006, Reich, 2014). We found that in the RIL population the genetic correlation between these traits was only 25% and each trait had three independent QTL. Thus divergence of these traits is likely due to independent loci which become structured across ecotypes through an accumulation of linkage disequilibrium resulting from strong directional or correlational selection. In this case, our crossing scheme was able to largely decouple these traits through recombination.

Observed pairwise epistatic interactions for root biomass, shoot biomass and root number showed that *hallii* alleles mask the effects of*filipes* alleles in all cases. When lines are homozygous for *hallii* parental alleles at CL5.3, it contributes to smaller phenotypes for these traits, regardless of the genotype at their respective interactive QTL. This suggests that the CL5.3 loci could include a pleiotropic gene with major effect that controls the development of multiple shoot and root size related traits. Natural populations of *P*. *hallii* ecotypes are largely homozygous, thus these linked QTL likely work together in a positive direction and contribute to the phenotypic trait correlations that underlie ecotype divergence. However, the observed epistasis in the RIL population could represent a Dobzhansky–Muller type incompatibility (Coyne & Orr, 2004; Sweigart & Willis, 2012; Bomblies, 2013), where these interactions in hybrid plants could be deleterious and impact survivorship by undermining synergistic trait relationships. The combination of reduced root and shoot size effected by *hallii* alleles is desirable in xeric environments, but maybe deleterious in the higher competition lowland environments *filipes* inhabits.

### Glasshouse detected genetic correlations confirmed under field conditions

There is persistent concern that effects observed in glasshouse studies are not representative of plant performance in natural or agronomic environments. Although glasshouses and growth chambers may be able to replicate a wide range of temperature and light conditions, other differences between these artificial and natural environments can be significant. Furthermore, glasshouse studies are often conducted on very young plants and in smaller than optimal pots, which can significantly alter root architectures compared to natural environments. Several recent studies have highlighted how differences in conditions between glasshouse and natural settings can affect the mapping of genetic architectures for various plant traits (Poorter *et.al*., 2012; reviewed in Lovell 2016).

We sought to overcome this concern by confirming the glasshouse detected genetic architecture of two of our chief traits of interest (root biomass and shoot biomass) in selected RIL lines and parental genotypes in a field setting at full plant maturity. In the RIL lines, we found that our glasshouse observed QTL were confirmed, even though the selected lines had differing genetic backgrounds. For the parental lines, we found that root mass ratio differences between the xeric and mesic ecotypes nearly doubled under field conditions as compared to the glasshouse study. This suggests that adaptive allocation of biomass to roots increases with plant age and can also be constrained by pot limitations in the glasshouse. More importantly, these results provide credence to the assumption that our glasshouse study is predictive of plant performance in a natural setting.

## Conclusion

In the process of ecotype formation, populations can diverge across many traits and exhibit different niche characteristics, which requires coordination between plant organ systems. Our study sheds light on the genetic architecture underlying the relationships between root and shoot traits involved in ecotype divergence of *Panicum hallii* and demonstrates that some correlated traits are under common genetic control as a result of QTL colocalization while other traits are controlled by independent loci. We found several genomic hotspots relating to multiple root and shoot traits and further insight into the molecular basis of these loci is an important step in understanding the genetic coordination of root and shoot systems involved in ecotype divergence. The RIL population utilized in this study has recently been increased to over 400 lines and should prove a valuable tool in investigating multiple facets of perennial grass biology.

## Acknowledgements

We would like to thank M. Donahue, J. Shih, L. Mayer, S. Faries, D. Miller and B. Campitelli for their help phenotyping in the glasshouse and field experiments; and J. Heiling, B. Whitaker and M. Stuke for help in generating RIL lines. Seeds for HAL2 and FIL2 parental lines were originally provided by the Lady Bird Johnson Wildflower Center and J. L. Reilly respectively; and the original cross of this material was created by D. Lowry. We also thank B. Campitelli and anonymous reviewers for discussion and comments that aided in the improvement of this manuscript. This research was supported by the DOE Office of Science, Office of Biological and Environmental Research (BER), grant no. DE-SC0008451 to T.E.J. Additional funding for this project came from an NSF Plant Genome Research Program Grant (IOS-0922457) to T.E.J and an NSF postdoctoral fellowship (IOS-1402393) to J.T.L. The work conducted by the U.S. Department of Energy Joint Genome Institute is supported by the Office of Science of the U.S. Department of Energy under Contract No. DE-AC02-05CH11231.

## Author Contributions

All authors contributed significantly to this work. A.K., J.T.L., J.E.B. and T.E.J. wrote the manuscript and contributed to statistical analysis. A.K. and T.E.J. designed experiments. A.K. and J.E.B. conducted experiments. J.J. and J.S. conducted the *Panicum hallii* genome assembly. Y.Y., J.J., and J.S. performed sequencing of the RIL population. J.T.L. created the genomic map.

## Supportive information Legends

**Table S1** Principal component loadings of measured traits in the *Panicum hallii* RIL population.

**Table S2** R/qtl input file for QTL mapping of measured traits in the *Panicum hallii* RIL population.

